# Hypothesis: Hypoxia induces *de novo* synthesis of NeuGc gangliosides in humans through CMAH domain substitute

**DOI:** 10.1101/125617

**Authors:** Paula A. Bousquet, Joe Alexander Sandvik, Nina F. Jeppesen Edin, Ute Krengel

## Abstract

Immunotherapy is a growing field in cancer research. A privileged tumor-associated antigen that has received much attention is *N*-glycolyl (NeuGc) GM3. This ganglioside is present in several types of cancer, but is almost undetectable in human healthy tissues. However, its non-hydroxylated variant, NeuAc GM3, is abundant in all mammals. Due to a deletion in the human gene encoding the key enzyme for synthesis of NeuGc, humans, in contrast to other mammals, cannot synthesize NeuGc GM3. Therefore the presence of this ganglioside in human cancer cells represents an enigma. It has been shown that hypoxic conditions trigger the expression of NeuGc gangliosides, which not only serve as attractive targets for cancer therapy, but also as diagnostic and prognostic tumor marker. Here, we confirm hypoxia-induced expression of the NeuGc GM3 ganglioside also in HeLa cells and reveal several candidate proteins, in particular GM3 synthase and subunit B of respiratory complex II (SDHB), that may be involved in the generation of NeuGc GM3 by SILAC-based proteome analysis. These findings have the potential to significantly advance our understanding of how this enigmatic tumor-associated antigen is produced in humans, and also suggest a possible mechanism of action of anti-tumor antibodies that recognize hypoxia markers, such as 14F7.

## Introduction

Cancer immunotherapy promises a revolution in cancer therapy, and was consequently selected by *Science* as Breakthrough of the year 2013 [1]. This type of therapy represents an entirely different approach to targeting and treating cancer compared to conventional therapies, such as radiation and chemotherapy. By selective targeting of tumor-specific antigens, treatment becomes more rational, with the potential for tailored personalized therapies.

One group of tumor-associated antigens with particularly attractive properties comprises sialic acid containing glycosphingolipids or gangliosides [2, 3]. In mammalian tissues, the most abundant forms of sialic acids are *N*-acetyl neuraminic acid (NeuAc) and *N*-glycolyl neuraminic acid (NeuGc), however, in human healthy tissue, NeuGc is present only in minute amounts, suggestive of contamination, rather than active generation [4, 5]. The synthesis of this glycolipid requires an enzyme, cytidine monophosphate-*N*-acetylneuraminic acid hydroxylase (CMAH), which catalyzes the hydroxylation of NeuAc to yield NeuGc. This enzyme is non-functional in humans due to a 92-bp deletion in the gene encoding its active site [6]. However, when healthy cells transform into malignant cells, NeuGc is expressed [7]. The origin of this change in carbohydrate profile is unknown. Approaches to solve this mystery have so far focused mainly on diet incorporation of NeuGc from animal sources [5, 8-11]. For example, it has recently been shown that *Cmah* knockout mice can incorporate NeuGc from dietary sources [12, 13]. NeuGc uptake and recycling by the lysosomal transporter sialin is well studied [14], and may be enhanced under hypoxic conditions [9]. Furthermore, an endogenous NeuGc-degrading pathway has been identified that produces *N*-acetylhexosamine metabolites, which may serve as precursors for the synthesis of synthesis of NeuGc [15, 16].

A NeuGc-containing ganglioside called NeuGc GM3 has attracted significant attention as tumor-associated antigen [7, 17, 18]. It is present on the surface of multiple types of cancers, such as breast cancer [19], melanoma [20], lung cancer [21] and retinoblastoma [22]. NeuGc GM3 not only has promising therapeutic and diagnostic potential but also holds a prognostic value, since its expression correlates with more aggressive disease [21, 23, 24]. Hypoxia has been recognized as an important and common characteristic of many advanced tumors, where progression occurs as a result of adaptation to and survival within the environment limited in oxygen [25]. In addition, hypoxia-induced cancer progression can affect the expression of cell surface antigens, and indeed, the expression of NeuGc gangliosides has been shown to be triggered by hypoxic condition [9]. Moreover, the aggressiveness of tumors correlates with *CMAH* expression levels [24], whereas in the brain, which is rich in oxygen, *CMAH* gene and NeuGc expression are suppressed [26]. However, the cellular mechanism causing the hypoxia-induced expression of NeuGc GM3 is still unknown.

NeuGc was originally considered an oncofetal antigen, since it is present in fetal tissue, but repressed during adult life. Normal embryonic development occurs in an environment low in oxygen, and hypoxia therefore also represents an important aspect of developmental morphogenesis [27, 28], promoting differentiation and proliferation [29, 30]. The fact that hypoxia promotes both tumor progression and embryonic development could suggest that alternative mechanisms to synthesize NeuGc gangliosides exist in humans, but that hypoxia is required to induce production. Indeed, this antigen has been detected in the epithelial lining of hollow organs and the endothelial lining of the vasculature [8, 10, 31]. A requirement for hypoxia for NeuGc expression would also explain why NeuGc is hardly detectable in the brains of most mammals, where the oxygen content is high [26], while other sialic acid variants are abundant in brain tissue [32].

Here, we used a combination of fluorescence-activated cell sorting (FACS) and stable isotope labeling by amino acids in cell culture (SILAC) to probe the origin of NeuGc GM3 expression, as triggered by hypoxia. SILAC is a quantitative proteomics method that reveals up- or down-regulated proteins upon exposure to different conditions [33]. The present work builds on a recent comprehensive SILAC study, where we analyzed protein regulation in HeLa cells in response to hypoxia [34]. Using an anti-tumor antibody that specifically recognizes NeuGc GM3 (14F7), we confirmed hypoxia-induced expression of this ganglioside in HeLa cells and identified several candidate proteins potentially involved in its *de novo* synthesis in humans, rescuing CMAH activity.

## Materials and Methods

### Cell cultivation

For hypoxic conditions, HeLa P cells were grown inside an InVivo2 400 incubator box (Ruskinn Technology, UK) where the oxygen level inside the box was set to 1%, while a standard CO_2_ incubator was used normoxic conditions. Prior to treatment, the cells were cultivated in Dulbecco’s modified Eagle’s medium (DMEM), supplemented with NeuGc free 10% human or chicken serum (H4522-Sigma, C5405-Sigma) and 1% penicillin/streptomycin for two to three weeks in order to ensure that no NeuGc was available in the medium. The experiment was performed in three biological replicates.

### Flow cytometry

After growing the cells for 72h in hypoxia or normoxia, they were trypsinized and thereafter washed twice with a solution containing 49 ml phosphate buffered saline (PBS), 1 ml of either fetal bovine serum, human or chicken serum, 200 µl EDTA before fixation with 4% paraformaldehyde for 15 minutes. After a third washing step, the cells were incubated in 50 µl of primary antibody (14F7, a murine anti-NeuGc GM3 antibody obtained from Center of Molecular Immunology, Havana) for 1h. The samples were then washed 2 times before 50µl of the secondary antibody was added (goat anti-mouse IgG1-FITC, Santa Cruz-2010) for 30 minutes in dark. After another washing step the cells were filtered and analysed by flow cytometry (BD Accuri C6 flow cytometer).

### Bioinformatic data analyses

Bioinformatics analysis was performed with STRING 9.1 (<u>http://string.embl.de/</u>) [35, 36], on data acquired previously by NanoLC-LTQ Orbitrap mass spectrometry, as described [34].

### Results and Discussion

Hypoxia-induced expression of NeuGc gangliosides has previously been observed in LS174T, Caco-2 and ZR-75-1 cells [9]. To probe if hypoxia also triggers increased expression of NeuGc GM3 in HeLa cells, flow cytometry experiments were performed. For detection, we used the murine antibody 14F7, which specifically recognizes NeuGc GM3 and can discriminate it from the very similar NeuAc derivative [20, 37-39]. This antibody was earlier subjected to glycoarray screening by the Functional Glycomics Consortium (www.functionalglycomics.org), with negative results, even for the NeuGc GM3 trisaccharide (data not shown), indicating that it is indeed very specific and furthermore requires the ganglioside’s sphingolipid part for antigen recognition. Since NeuGc GM3 is abundant in most mammalian tissues (except brain tissue), but deficient in healthy human adults [26, 40], we cultivated the cells in human serum for 2-3 weeks before analysis. Figure 1 shows that hypoxic conditions indeed induced a three-fold increase in expression of NeuGc GM3 in HeLa cells, in support of the hypothesis that hypoxia is an important factor for the generation of this antigen. To confirm our results, we repeated the experiment using chicken serum. Chicken, like other birds (as well as humans, reptiles and platypus), are deficient in NeuGc [41]. We observed that hypoxia induced a similar two-fold increase in expression of NeuGc GM3 in chicken serum, as determined by FACS analysis with 14F7 (Figure S1), whereas culturing in NeuGc-rich fetal bovine serum showed no such effect (Figure S2), in keeping with the hypothesis that *de novo* NeuGc synthesis can be induced by hypoxia.

**Figure 1.**
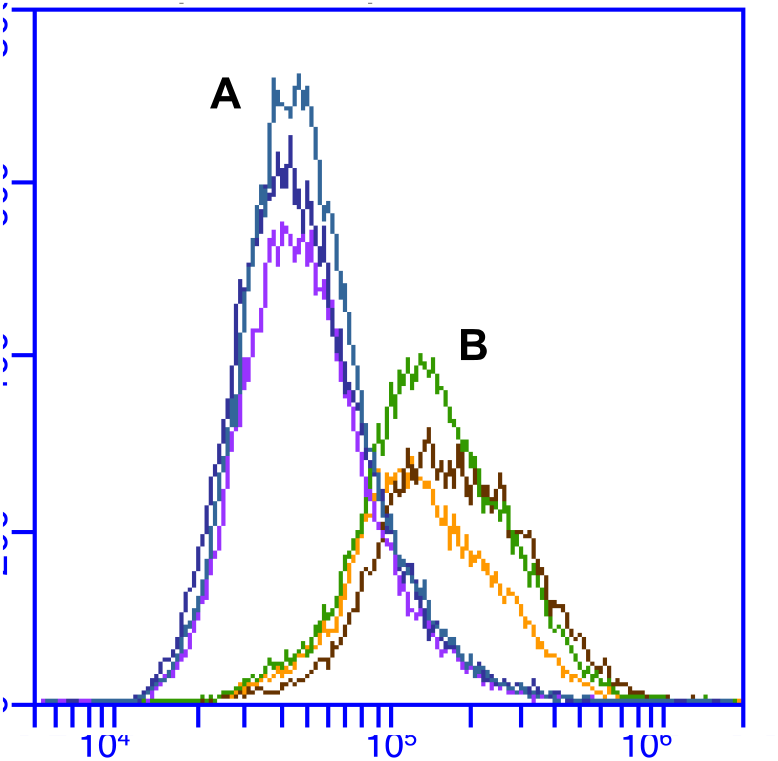
NeuGc GM3 expression in normoxia (A) and hypoxia (B). Using human serum, the tumor antigen showed three-fold increased expression under hypoxic conditions. Experiments were performed in biological triplicates, and the figure shows an overlay of the triplicate samples.

But how does hypoxia trigger the expression of NeuGc GM3? What are the cellular mechanisms involved, and is *de novo* synthesis required or could other factors be at play? It has been suggested that hypoxic culture induces the expression of sialin, a sialic acid transporter, leading to increased incorporation of NeuGc from the external environment [9]. Indeed, a recent study of aggressive melanoma showed that the amount of NeuGc GM3 depended on the external medium [24]. However, we did not observe an increased expression of NeuGc GM3 using media supplemented with fetal bovine serum, which contains NeuGc (Figure S2). Sialin (SLC17A5) was not identified in our recent SILAC study, not even at below-significance levels, even though 26 other proteins from the SCL-family were detected, of which four were up-regulated [34]. One of them was the GM3 synthase (ST3GAL5/SLC35E1) responsible for catalyzing the covalent addition of sialic acid to lactosylceramide to generate the GM3 ganglioside. Upregulation of GM3 synthase (1.5 fold; p = 0.11) was below statistical significance, but was confirmed by Western blot analysis (Figure 2).

**Figure 2.**
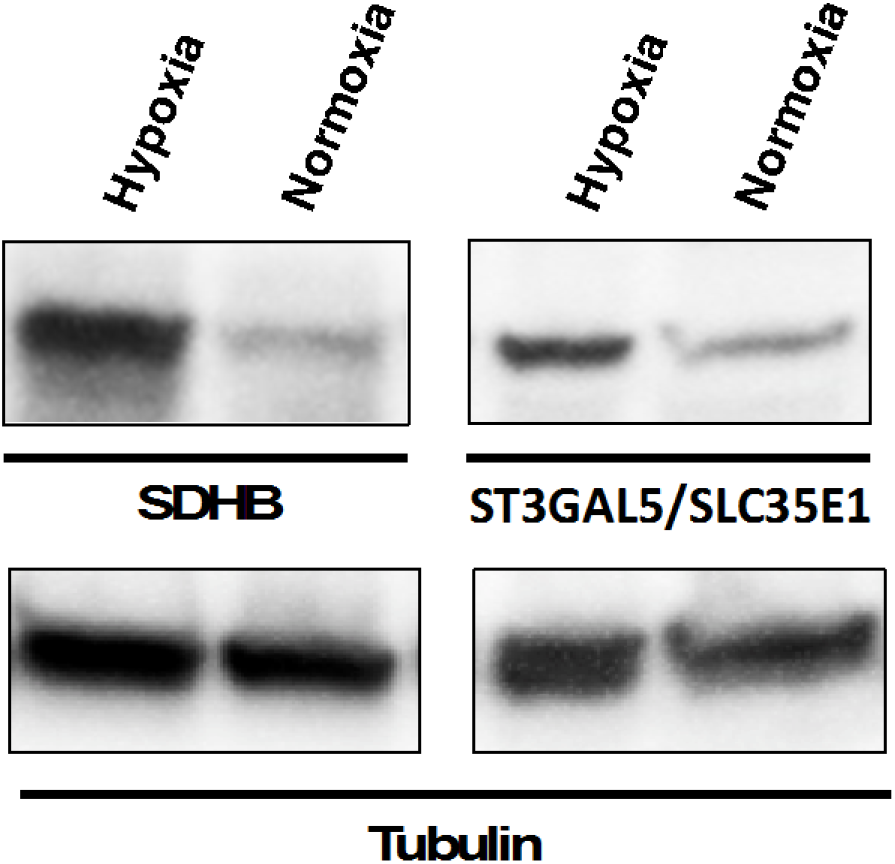
Western blot analysis of SDHB and GM3 synthase. Representative Western blots for succinate dehydrogenase subunit B and ST3GAL5/SLC35E1 from HeLa cells grown are shown, comparing hypoxic or normoxic conditions. The experiments were performed in biological triplicates. Additionally, the expression of β-tubulin is shown to demonstrate an equal loading of gels.

In the same study, we found an iron-sulfur [Fe_2_S_2_]-containing protein to be significantly up-regulated under hypoxic conditions (1.9 fold; p=0.0057), while many other mitochondrial proteins were down-regulated [34]. The protein in question is subunit B of the succinate dehydrogenase complex (which represents complex II of the respiratory chain). It is only this subunit (SDHB) that was found up-regulated, while the other subunits of complex II were not affected. We have now confirmed hypoxia-induced up-regulation of SDHB by Western blot analysis (Figure 2). This is interesting since the *cmah* gene responsible for NeuGc synthesis in mammals exhibits a 92-bp deletion in humans. The deleted gene fragment codes for a Rieske domain, an [Fe_2_S_2_]-cluster, which presumably comprises the active site of the enzyme. – Could it be that SDHB can take on the function of the missing Rieske domain? Or do other mechanisms lie at the heart of hypoxia-induced NeuGc GM3 expression?

Gangliosides can rearrange in the membrane and create “clusters” together with cholesterol (referred to as ‘lipid rafts’), forming more ordered regions of the membrane [42-44]. The density, presentation and organization of glycosphingolipids may affect antibody specificity [3, 45, 46], and it has been shown that some ganglioside antigens are not fully antigenic and thus fail to interact with antibodies when their density is below a threshold value [47]. Currently, we cannot exclude that the increased binding of the 14F7 antibody after hypoxia treatment is caused or enhanced by such membrane remodeling. Another possibility is the recycling of gangliosides from other compartments in the cell and transport to the cell surface, where antibody detection occurs.

Nevertheless, it is intriguing that GM3 synthase appears to be up-regulated under hypoxic conditions, suggesting that *in vivo* NeuGc GM3 synthesis might actually occur. This would also explain the enhanced *CMAH* expression levels observed in aggressive human tumors [24] and the low expression levels in the vertebrate brain [26], protecting the brain from NeuGc’s (or CMAH’s) presumed toxic effects. If this can be confirmed, we may have discovered a salvaging pathway for the biosynthesis of NeuGc that is induced by hypoxia. The central player of our hypothesis is subunit B from respiratory complex II, or SDHB. Figure 3 shows how SDHB is clearly associated with mitochondrial deficiency, affecting clusters of mitochondrial ribosomal proteins (MRPs) and translocases of the inner and outer mitochondrial membrane (TIMM/TOMM) (down-regulated), while glycolysis is up-regulated. Whether this iron-sulfur domain could, provided hypoxic conditions, replace the function of the original Rieske domain is yet to be investigated. Mitochondria are the major consumers of iron in the cell and hence the primary sites for the synthesis of iron-sulfur clusters [48]. Since Fe/S clusters are required for viability and exist in all kingdoms of life, the loss of these components is lethal during early embryonic development, and mutations in genes encoding proteins with such functionalities can cause severe disease in humans [48, 49]. Similarly, Fe/S clusters may be important for survival and growth of cancer cells, by inducing NeuGc ganglioside synthesis.

**Figure 3.**
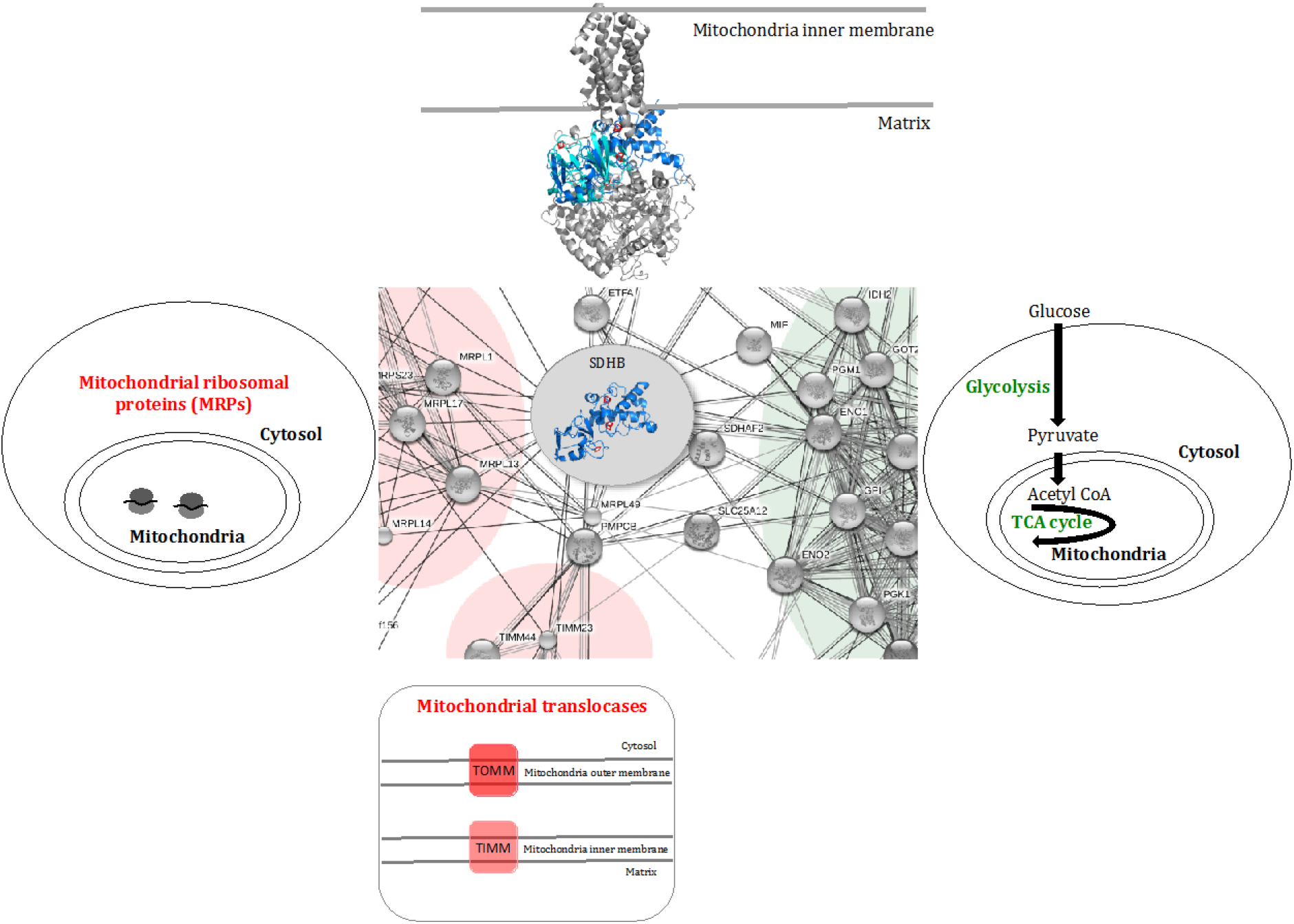
Succinate dehydrogenase subunit B (blue) and associations to up- or down-regulated clusters in response to hypoxia. The STRING bioinformatical tool (<u>http://string.embl.de/</u>) was used to visualize known and predicted protein-protein interactions. The 3D structure of the entire succinate dehydrogenase complex is shown on top, with subunit B (blue) aligned with the Rieske domain (turquoise). Iron-sulfur clusters are depicted in red.

Another interesting aspect of this study relates to cancer immunotherapy. The strong evidence for hypoxia causing tumor progression and its importance in therapy resistance makes hypoxia-regulated molecules valuable therapeutic targets. Anti-tumor antibodies that recognize or mimic such hypoxia markers, like 14F7 and 1E10 (racotumumab), may owe their promising clinical potential [17, 50-56] at least in part to their ability to home in on the tumor center or generate anti-idiotypic antibodies with this ability. Inside the tumor, where hypoxia rules, expression of the targets is greatest; hence antibody binding will be strongest. Initial binding to the tumor surface should hence trigger the antibody’s walk to the tumor center, to treat it from its roots. Antibodies recognizing hypoxia-rich regions of the tumor, especially with highly dynamic association-dissociation kinetics, should hence be highly attractive for cancer therapies, either stand-alone or as potential adjuvants.

## Acknowledgments

We wish to thank Prof. Erik O. Pettersen (Physics Department, University of Oslo) and Dr. Espen Stang (Oslo University Hospital) for access to their laboratories and support, and Vibeke Bertelsen and Marianne Skeie Rødland for practical tips. We further acknowledge the Center for Molecular Immunology in Havana for providing us with the antibody 14F7 and the Consortium for Functional Glycomics for glycan array analysis. The HeLa cell line was derived from Henrietta Lacks in 1951, who made significant contributions to biomedical research. We would like to thank her and her family members for this.

## Abbreviations

CMAH: Cytidine monophosphate-*N*-acetylneuraminic acid hydroxylase
DMEM: Dulbecco’s modified Eagle’s medium; ethylenediaminetetraacetic acid
FACS: Fluorescence-activated cell sorting
FITC: Fluorescein isothiocyanate
MRP: Mitochondrial ribosomal protein
NeuAc: *N*-acetyl neuraminic acid
NeuGc: *N*-glycolyl neuraminic acid
SDHB: Succinate dehydrogenase complex subunit B
SILAC: Stable isotope labeling by amino acid in cell culture
TIMM: Translocases of the inner mitochondrial membrane
TOMM: Translocases of the outer mitochondrial membrane
PBS: Phosphate buffered saline
LC-LTQ: Liquid chromatography – linear trap quadrupole.

## Supplementary Figures

**Figure S1.**
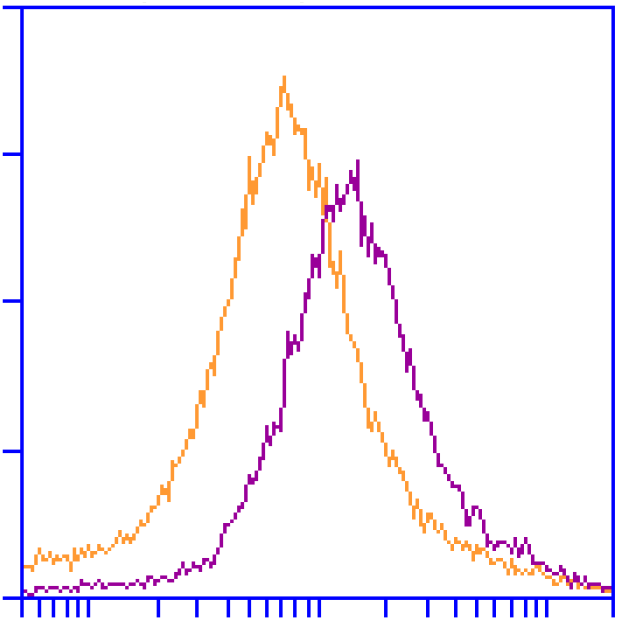
Flow cytometry histogram showing NeuGc GM3 expression in HeLa cells grown in media supplemented with chicken serum under normoxic (yellow) and hypoxic (purple) conditions. As for the experiments with human serum, the tumor antigen showed increased expression under hypoxic conditions (here: two-fold increase). Experiments were performed in biological triplicates, with reproducible results. One representative histogram is shown in the figure.

**Figure S2.**
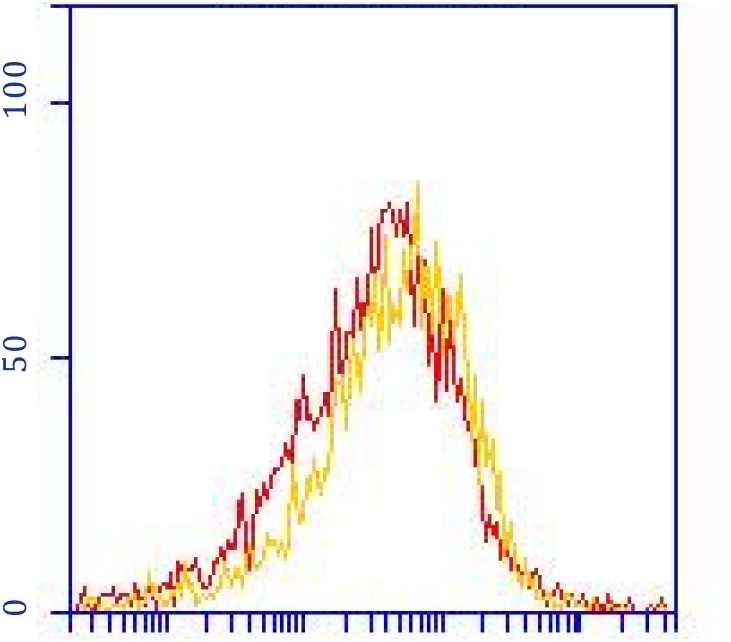
Flow cytometry histogram showing NeuGc GM3 expression in HeLa cells grown in media supplemented with fetal bovine serum under normoxic (red) and hypoxic (yellow) conditions. In contrast to the experiments with human and chicken serum, the tumor antigen did not show increased expression under hypoxic conditions. Experiments were performed in biological triplicates, with reproducible results. One representative histogram is shown in the figure.

## References

[1] J. Couzin-Frankel, Breakthrough of the year 2013. Cancer immunotherapy, Science, 342 (2013) 1432–1433.

[2] M.A. Cheever, J.P. Allison, A.S. Ferris, O.J. Finn, B.M. Hastings, T.T. Hecht, I. Mellman, S.A. Prindiville, J.L. Viner, L.M. Weiner, L.M. Matrisian, The prioritization of cancer antigens: a national cancer institute pilot project for the acceleration of translational research, Clin Cancer Res, 15 (2009) 5323–5337.

[3] U. Krengel, P.A. Bousquet, Molecular recognition of gangliosides and their potential for cancer immunotherapies, Front Immunol, 5 (2014) 325.

[4] A. Irie, S. Koyama, Y. Kozutsumi, T. Kawasaki, A. Suzuki, The molecular basis for the absence of *N*-glycolylneuraminic acid in humans, J Biol Chem, 273 (1998) 15866–15871.

[5] A. Varki, *N*-glycolylneuraminic acid deficiency in humans, Biochimie, 83 (2001) 615–622.

[6] H.H. Chou, H. Takematsu, S. Diaz, J. Iber, E. Nickerson, K.L. Wright, E.A. Muchmore, D.L. Nelson, S.T. Warren, A. Varki, A mutation in human CMP-sialic acid hydroxylase occurred after the *Homo-Pan* divergence, Proc Natl Acad Sci USA, 95 (1998) 11751–11756.

[7] Y.N. Malykh, R. Schauer, L. Shaw, *N*-glycolylneuraminic acid in human tumours, Biochimie, 83 (2001) 623–634.

[8] P. Tangvoranuntakul, P. Gagneux, S. Diaz, M. Bardor, N. Varki, A. Varki, E. Muchmore, Human uptake and incorporation of an immunogenic nonhuman dietary sialic acid, Proc Natl Acad Sci U S A, 100 (2003) 12045–12050.

[9] J. Yin, A. Hashimoto, M. Izawa, K. Miyazaki, G.-Y. Chen, H. Takematsu, Y. Kozutsumi, A. Suzuki, K. Furuhata, F.-L. Cheng, C.-H. Lin, C. Sato, K. Kitajima, R. Kannagi, Hypoxic culture induces expression of sialin, a sialic acid transporter, and cancer-associated gangliosides containing non-human sialic acid on human cancer cells, Cancer Res, 66 (2006) 2937–2945.

[10] E. Byres, A.W. Paton, J.C. Paton, J.C. Löfling, D.F. Smith, M.C. Wilce, U.M. Talbot, D.C. Chong, H. Yu, S. Huang, X. Chen, N.M. Varki, A. Varki, J. Rossjohn, T. Beddoe, Incorporation of a non-human glycan mediates human susceptibility to a bacterial toxin, Nature, 456 (2008) 648–652.

[11] F. Alisson-Silva, K. Kawanishi, A. Varki, Human risk of diseases associated with red meat intake: Analysis of current theories and proposed role for metabolic incorporation of a non-human sialic acid, Mol Aspects Med, 51 (2016) 16–30.

[12] K. Banda, C.J. Gregg, R. Chow, N.M. Varki, A. Varki, Metabolism of vertebrate amino sugars with *N*-glycolyl groups: mechanisms underlying gastrointestinal incorporation of the non-human sialic acid xeno-auto-antigen *N*-glycolylneuraminic acid, J Biol Chem, 287 (2012) 28852–28864.

[13] A.K. Bergfeld, R. Lawrence, S.L. Diaz, O.M.T. Pearce, D. Ghaderi, P. Gagneux, M.G. Leakey, A. Varki, *N*-glycolyl groups of nonhuman chondroitin sulfates survive in ancient fossils, Proc Natl Acad Sci U S A, 114 (2017) E8155–E8164.

[14] M. Bardor, D.H. Nguyen, S. Diaz, A. Varki, Mechanism of uptake and incorporation of the non-human sialic acid *N*-glycolylneuraminic acid into human cells, J Biol Chem, 280 (2005) 4228–4237.

[15] A.K. Bergfeld, O.M. Pearce, S.L. Diaz, T. Pham, A. Varki, Metabolism of vertebrate amino sugars with *N*-glycolyl groups: elucidating the intracellular fate of the non-human sialic acid *N*-glycolylneuraminic acid, J Biol Chem, 287 (2012) 28865–28881.

[16] A.K. Bergfeld, O.M. Pearce, S.L. Diaz, R. Lawrence, D.J. Vocadlo, B. Choudhury, J.D. Esko, A. Varki, Metabolism of vertebrate amino sugars with *N*-glycolyl groups: incorporation of *N*-glycolylhexosamines into mammalian glycans by feeding *N*-glycolylgalactosamine, J Biol Chem, 287 (2012) 28898–28916.

[17] L.E. Fernandez, M.R. Gabri, M.D. Guthmann, R.E. Gomez, S. Gold, L. Fainboim, D.E. Gomez, D.F. Alonso, NGcGM3 ganglioside: a privileged target for cancer vaccines, Clin Dev Immunol, 2010 (2010) 814397.

[18] A.N. Samraj, H. Läubli, N. Varki, A. Varki, Involvement of a non-human sialic acid in human cancer, Front Oncol, 4 (2014) 33.

[19] G. Marquina, H. Waki, L.E. Fernandez, K. Kon, A. Carr, O. Valiente, R. Perez, S. Ando, Gangliosides expressed in human breast cancer, Cancer Res, 56 (1996) 5165–5171.

[20] A. Carr, A. Mullet, Z. Mazorra, A.M. Vázquez, M. Alfonso, C. Mesa, E. Rengifo, R. Pérez, L.E. Fernández, A mouse IgG1 monoclonal antibody specific for *N*-glycolyl GM3 ganglioside recognized breast and melanoma tumors, Hybridoma, 19 (2000) 241–247.

[21] R. Blanco, C.E. Rengifo, M. Cedeño, M. Frómeta, E. Rengifo, A. Carr, Immunoreactivity of the 14F7 Mab (raised against *N*-glycolyl GM3 ganglioside) as a positive prognostic factor in non-small-cell lung cancer, Patholog Res Int, 2012 (2012) 235418.

[22] A.V. Torbidoni, A. Scursoni, S. Camarero, V. Segatori, M. Gabri, D. Alonso, G. Chantada, M.T. de Dávila, Immunoreactivity of the 14F7 Mab raised against *N*-glycolyl GM3 ganglioside in retinoblastoma tumours, Acta Ophthalmol, 93 (2015) e294–300.

[23] A.M. Scursoni, L. Galluzzo, S. Camarero, J. Lopez, F. Lubieniecki, C. Sampor, V.I. Segatori, M.R. Gabri, D.F. Alonso, G. Chantada, M.T. de Dávila, Detection of *N*-glycolyl GM3 ganglioside in neuroectodermal tumors by immunohistochemistry: an attractive vaccine target for aggressive pediatric cancer, Clin Dev Immunol, 2011 (2011) 245181.

[24] C. Tringali, I. Silvestri, F. Testa, P. Baldassari, L. Anastasia, R. Mortarini, A. Anichini, A. López-Requena, G. Tettamanti, B. Venerando, Molecular subtyping of metastatic melanoma based on cell ganglioside metabolism profiles, BMC Cancer, 14 (2014) 560.

[25] K. DeClerck, R.C. Elble, The role of hypoxia and acidosis in promoting metastasis and resistance to chemotherapy, Front Biosci (Landmark Ed), 15 (2010) 213–225.

[26] L.R. Davies, A. Varki, Why Is *N*-glycolylneuraminic acid rare in the vertebrate brain?, Top Curr Chem, 366 (2015) 31–54.

[27] G.J. Burton, E. Jauniaux, A.L. Watson, Maternal arterial connections to the placental intervillous space during the first trimester of human pregnancy: the Boyd collection revisited, Am J Obstet Gynecol, 181 (1999) 718–724.

[28] S.L. Dunwoodie, The role of hypoxia in development of the mammalian embryo, Dev Cell, 17 (2009) 755–773.

[29] C.E. Forristal, K.L. Wright, N.A. Hanley, R.O. Oreffo, F.D. Houghton, Hypoxia inducible factors regulate pluripotency and proliferation in human embryonic stem cells cultured at reduced oxygen tensions, Reproduction, 139 (2010) 85–97.

[30] S. Prado-Lopez, A. Conesa, A. Armiñán, M. Martínez-Losa, C. Escobedo-Lucea, C. Gandia, S. Tarazona, D. Melguizo, D. Blesa, D. Montaner, S. Sanz-González, P. Sepúlveda, S. Götz, J.E. O'Connor, R. Moreno, J. Dopazo, D.J. Burks, M. Stojkovic, Hypoxia promotes efficient differentiation of human embryonic stem cells to functional endothelium, Stem Cells, 28 (2010) 407–418.

[31] T. Pham, C.J. Gregg, F. Karp, R. Chow, V. Padler-Karavani, H. Cao, X. Chen, J.L. Witztum, N.M. Varki, A. Varki, Evidence for a novel human-specific xeno-auto-antibody response against vascular endothelium, Blood, 114 (2009) 5225–5235.

[32] M. Iwamori, Y. Nagai, A new chromatographic approach to the resolution of individual gangliosides. Ganglioside mapping, Biochim Biophys Acta, 528 (1978) 257–267.

[33] S.E. Ong, B. Blagoev, I. Kratchmarova, D.B. Kristensen, H. Steen, A. Pandey, M. Mann, Stable isotope labeling by amino acids in cell culture, SILAC, as a simple and accurate approach to expression proteomics, Mol Cell Proteomics, 1 (2002) 376–386.

[34] P.A. Bousquet, J.A. Sandvik, M.Ø. Arntzen, N.F. Jeppesen Edin, S. Christoffersen, U. Krengel, E.O. Pettersen, B. Thiede, Hypoxia strongly affects mitochondrial ribosomal proteins and translocases, as shown by quantitative proteomics of HeLa cells, Int J Proteomics, 2015 (2015) 678527.

[35] B. Snel, G. Lehmann, P. Bork, M.A. Huynen, STRING: a web-server to retrieve and display the repeatedly occurring neighbourhood of a gene, Nucleic Acids Res, 28 (2000) 3442–3444.

[36] A. Franceschini, D. Szklarczyk, S. Frankild, M. Kuhn, M. Simonovic, A. Roth, J. Lin, P. Minguez, P. Bork, C. von Mering, L.J. Jensen, STRING v9.1: protein-protein interaction networks, with increased coverage and integration, Nucleic Acids Res, 41 (2013) D808–815.

[37] G. Rojas, A. Talavera, Y. Munoz, E. Rengifo, U. Krengel, J. Angström, J. Gavilondo, E. Moreno, Light-chain shuffling results in successful phage display selection of functional prokaryotic-expressed antibody fragments to *N*-glycolyl GM3 ganglioside, J Immunol Methods, 293 (2004) 71–83.

[38] U. Krengel, L.-L. Olsson, C. Martinez, A. Talavera, G. Rojas, E. Mier, J. Angström, E. Moreno, Structure and molecular interactions of a unique antitumor antibody specific for *N*-glycolyl GM3, J Biol Chem, 279 (2004) 5597–5603.

[39] G. Rojas, A. Pupo, S. Gómez, U. Krengel, E. Moreno, Engineering the binding site of an antibody against *N*-glycolyl GM3: from functional mapping to novel anti-ganglioside specificities, ACS Chem Biol, 8 (2013) 376–386.

[40] S. Hara, Y. Takemori, M. Yamaguchi, M. Nakamura, Y. Ohkura, Fluorometric high-performance liquid chromatography of *N*-acetyl- and *N*-glycolylneuraminic acids and its application to their microdetermination in human and animal sera, glycoproteins, and glycolipids, Anal Biochem, 164 (1987) 138–145.

[41] R. Schauer, G.V. Srinivasan, B. Coddeville, J.P. Zanetta, Y. Guérardel, Low incidence of *N*-glycolylneuraminic acid in birds and reptiles and its absence in the platypus, Carbohydr Res, 344 (2009) 1494–1500.

[42] K. Simons, E. Ikonen, Functional rafts in cell membranes, Nature, 387 (1997) 569–572.

[43] K. Simons, D. Toomre, Lipid rafts and signal transduction, Nat Rev Mol Cell Biol, 1 (2000) 31–39.

[44] K. Simons, J.L. Sampaio, Membrane organization and lipid rafts, Cold Spring Harb Perspect Biol, 3 (2011) a004697.

[45] M.L. DeMarco, R.J. Woods, Atomic-resolution conformational analysis of the GM3 ganglioside in a lipid bilayer and its implications for ganglioside-protein recognition at membrane surfaces, Glycobiology, 19 (2009) 344–355.

[46] T. Róg, I. Vattulainen, Cholesterol, sphingolipids, and glycolipids: What do we know about their role in raft-like membranes?, Chem Phys Lipids, 184 (2014) 82–104.

[47] G.A. Nores, T. Dohi, M. Taniguchi, S. Hakomori, Density-dependent recognition of cell surface GM3 by a certain anti-melanoma antibody, and GM3 lactone as a possible immunogen: requirements for tumor-associated antigen and immunogen, J Immunol, 139 (1987) 3171–3176.

[48] O. Stehling, R. Lill, The role of mitochondria in cellular iron-sulfur protein biogenesis: mechanisms, connected processes, and diseases, Cold Spring Harb Perspect Biol, 5 (2013) a011312.

[49] M. Cossée, H. Puccio, A. Gansmuller, H. Koutnikova, A. Dierich, M. LeMeur, K. Fischbeck, P. Dollé, M. Koenig, Inactivation of the Friedreich ataxia mouse gene leads to early embryonic lethality without iron accumulation, Hum Mol Genet, 9 (2000) 1219–1226.

[50] A. Carr, C. Mesa, M. del Carmen Arango, A.M. Vázquez, L.E. Fernández, *In vivo* and *in vitro* anti-tumor effect of 14F7 monoclonal antibody, Hybrid Hybridomics, 21 (2002) 463–468.

[51] M. Alfonso, A. Díaz, A.M. Hernández, A. Pérez, E. Rodríguez, R. Bitton, R. Pérez, A.M. Vázquez, An anti-idiotype vaccine elicits a specific response to *N*-glycolyl sialic acid residues of glycoconjugates in melanoma patients, J Immunol, 168 (2002) 2523–2529.

[52] A. Carr, E. Rodríguez, C. Arango Mdel, R. Camacho, M. Osorio, M. Gabri, G. Carrillo, Z. Valdés, Y. Bebelagua, R. Pérez, L.E. Fernández, Immunotherapy of advanced breast cancer with a heterophilic ganglioside (NeuGcGM3) cancer vaccine, J Clin Oncol, 21 (2003) 1015–1021.

[53] A.M. Hernández, D. Toledo, D. Martínez, T. Griñán, V. Brito, A. Macías, S. Alfonso, T. Rondón, E. Suárez, A.M. Vázquez, R. Pérez, Characterization of the antibody response against NeuGcGM3 ganglioside elicited in non-small cell lung cancer patients immunized with an anti-idiotype antibody, J Immunol, 181 (2008) 6625–6634.

[54] Z. Gajdosik, Racotumomab - a novel anti-idiotype monoclonal antibody vaccine for the treatment of cancer, Drugs Today (Barc), 50 (2014) 301–307.

[55] S. Alfonso, A. Valdés-Zayas, E.R. Santiesteban, Y.I. Flores, F. Areces, M. Hernández, C.E. Viada, I.C. Mendoza, P.P. Guerra, E. García, R.A. Ortiz, A.V. de la Torre, M. Cepeda, K. Pérez, E. Chong, A.M. Hernández, D. Toledo, Z. González, Z. Mazorra, T. Crombet, R. Pérez, A.M. Vázquez, A.E. Macías, A randomized, multicenter, placebo-controlled clinical trial of racotumomab-alum vaccine as switch maintenance therapy in advanced non-small cell lung cancer patients, Clin Cancer Res, 20 (2014) 3660–3671.

[56] J.M. Brown, A.J. Giaccia, The unique physiology of solid tumors: opportunities (and problems) for cancer therapy, Cancer Res, 58 (1998) 1408–1416.

